# RNA Strain-Match: A tool for matching single-nucleus, single-cell, or bulk RNA-sequencing alignment data to its corresponding genotype

**DOI:** 10.1101/2023.07.14.548847

**Authors:** Jon A. L. Willcox, Maria A. Telpoukhovskaia, Niran Hadad, Stephanie M. Boas, Amy Dunn, Michael C. Saul, David G. Ashbrook, Robert W. Williams, Kristen M. S. O’Connell, Catherine C. Kaczorowski

## Abstract

When next generation sequencing is performed in large batches, there are several stages at which samples can be swapped or mislabeled. It is therefore helpful, when possible, to integrate measures into analysis pipelines to confirm that samples match their assigned metadata. Here, we introduce RNA Strain-Match (GitHub: https://github.com/jon-willcox/RNA-strain-match), a quality control tool developed to match RNA data in the form of sequence alignment files (*i.e*. SAM or BAM files) to their corresponding genotype without the use of an RNA variant call format file. We successfully used RNA Strain-Match in tandem with assessment of markers for sex and transgene status to identify and correct sample mismatches in 50/379 samples (13%) from two distinct recombinant inbred mouse models (BXD and Collaborative Cross). We believe this tool will be beneficial to any research group working with similar data.

## Introduction

For large batches of next generation sequencing data, it can be helpful to include quality control (QC) steps to ensure that all samples are correctly labelled and assigned their corresponding metadata. These steps may include verifying sample sex via identification of sex-specific chromosomal markers or confirming the presence of genomic features known to vary across samples, *e.g*. transgenic insertions. Perhaps the most reliable method of distinguishing between samples, however, is to compare the genomic sequence to a reference. This can be done in a variety of ways, depending on the study and the available resources; if sample-specific genotyping is available, a direct comparison is ideal; however, establishing relatedness to familial or genetically similar samples can also be an effective approach.

RNA Strain-Match uses known genotyping information – specifically autosomal coding single nucleotide polymorphisms (SNPs) with a single alternative allele – to match RNA sequencing data to corresponding genotypic information. It is designed to accept alignment data in the form of BAM files as input, which is convenient as these files are often generated as an intermediate step during nuclear, single-cell, and bulk RNA-seq processing. For example, the default settings for commonly used aligners such as STAR and Cell Ranger produce BAM files prior to calculating transcript counts. If these files are available, they can be used directly as input to RNA Strain-Match without additional formatting steps.

RNA Strain-Match was originally developed to check assignments of single-nucleus RNA sequencing (NucSeq) data to samples in the AD-BXD panel,^1^ which comprise of F1 progeny from B6.Cg-5XFAD hemizygous transgenic dams crossed with various strains from the BXD (B6 x DBA/2J (D2)) genetic reference panel recombinant inbred mouse lines^2^ (studies 1, 2, 4-6 in Table 1), a dataset that introduces challenges that are not present when genotyping samples from genetically unique individuals. Because each variant in recombinant inbred mice is shared across approximately 50% of strains, it is particularly important to genotype from SNPs widely distributed across the genome to ensure that sufficient recombinant regions are sampled for reliable strain identification. Additionally, all sites are expected to be either homozygous for the *B*-allele (*B*/*B*) or heterozygous (*B*/*D*), where *B* or *D* is the allele from the B6 or D2 mouse, respectively. This means that the presence of a single read with a *D*-allele should indicate heterozygosity. However, because NucSeq is frequently contaminated by small amounts of cell-free RNA reads derived from unrelated samples, it is possible for contaminant *D*-alleles to be present in small proportions. We therefore incorporated options for read-depth and ALT-allele fraction cutoffs to ensure that matching scores are robust against low-level contamination. While we have tested it with mouse data herein, this tool should be applicable to any RNA data if the appropriate genotyping data are available.

**Table 1.** Summary table of six studies included in trial analysis, including the number of samples, the tissue type, the genome the data was aligned to, and the genetic panel of the samples.

| Study | Number of Samples | Tissue | Genome | Panel |
| --- | --- | --- | --- | --- |
| 1 | 56 | Hippocampus (HC) | mm10 | AD-BXD |
| 2 | 56 | Frontal Cortex (Ctx) | mm10 | AD-BXD |
| 3 | 85 | Hippocampus (HC) | mm10 | CC |
| 4 | 18 | Hippocampus (HC) | mm10 | AD-BXD |
| 5 | 159 | Frontal Cortex (Ctx) | mm39+5XFAD | AD-BXD |
| 6 | 5 | Hypothalamus (HT) | mm39+5XFAD | AD-BXD |

## Methods

### RNA Strain-Match Method

The full RNA Strain-Match pipeline can be found on GitHub at https://github.com/jon-willcox/RNA-strain-match. Strain genotyping information is provided by the user in the form of a tab-delimited matrix specifying the genotype at autosomal, coding SNPs with a single alternative allele (one per row) for each strain, where “0/0” is homozygous reference, “1/1” is homozygous alternative, “0/1” is heterozygous. The first six columns in this file (“X.CHROM”, “POS”, “ID”, “REF”, “ALT”, “QUAL”) match the first six columns of a variant call format (VCF) file, and are followed by a column for each strain and a column (“VAR”) with a unique variant ID (a recommended procedure for building a new genotype file can be found on the GitHub page).

RNA-sequencing data is provided as a binary alignment map (BAM) file. The samtools mpileup function is used to extract all alleles present at each SNP location.^3,4^ Non-ref/alt alleles are discarded and only positions with sufficient depth of coverage (assigned via the “dp_cut” variable) are included in downstream analysis. At each position, the alternative allele fraction is calculated as N_ALT_/(N_ALT_ + N_REF_), where N_ALT_ and N_REF_ are the number of reads containing the alternative and reference alleles, respectively. Each position is then characterized by the absence or presence of an alternative allele (REF and ALT, respectively). A position is called as REF if the alternative allele is entirely absent, whereas it is considered to be ALT (either homozygous or heterozygous) if the alternative allele fraction (AF_ALT_) is greater than a user-specified cutoff (assigned via the “af_cut” variable). For F1 progeny, REF calls are considered to match 0/0 or 0/1 genotyping, and ALT calls are considered to match 0/1 or 1/1 genotyping. For non-F1 samples, the same matching criteria is used except REF calls are only considered to match 0/0 genotyping. The AF_ALT_ cutoff makes matching robust in the presence of contamination, which can occur, for example, in the form of cell-free RNA in NucSeq data.^5^

Once all SNPs are evaluated as REF or ALT, a score is assigned to the sample for each strain as the percent of informative SNPs (*i.e*. all SNPs with a REF/ALT assignment) that match the strain.

### Trial Dataset

RNA Strain-Match was used to confirm the strain of 294 hippocampus (HC), hypothalamus (HT), and frontal cortex (Ctx) NucSeq samples from F1 progeny of B6 congenic 5XFAD hemizygous transgenic females (JAX stock #34848)^6^ crossed with B6 (C57BL/6J; JAX stock #000664), D2Gpnmb (DBA/2J-*Gpnmb*^*+*^/SjJ; JAX stock 007048)^7^, or one of 23 BXD strains (BXD14, BXD19, BXD28, BXD33, BXD38, BXD39, BXD50, BXD53, BXD61, BXD65, BXD66, BXD73, BXD74, BXD75, BXD77, BXD83, BXD99, BXD112, BXD113, BXD124, or BXD161)^8^, (*i.e*., the AD-BXD panel)^1^ as well as 85 HC samples from the Collaborative Cross (CC) family of recombinant inbred mice from eight founder strains (CC003, CC004, CC005, CC006, CC019, CC032, CC041, and CC061).^9^ More specifically, samples were collected from HC (studies 1, 3, and 4; N=159), Ctx (studies 2 and 5; N=215), and HT (study 6; N=5) (Tables 1 and S1). RNA extraction and sequencing for all samples was performed as described previously by Telpoukhovskaia, et al.^10^

### Sequencing Alignment

NucSeq data were aligned using the CellRanger count pipeline V3.1.0 (10x Genomics) to either the GRCm38 genome build (mm10; n=215) or the GRCm39 build including the 5XFAD transgene^6^ (mm39+5XFAD; n=164). For all alignments, the reference genome (GRCm38 or GRCm39) is derived from the B6 mouse and the *B*-allele is the reference allele.

### Match Calculation

Matching scores for each B6xBXD sample were calculated using 114,977 autosomal coding SNPs that differ between B6 and D2.^11^ In CC samples, scores were generated using 874,953 autosomal coding SNPs^9,12^ that differ among the 8 CC founder strains. Of these SNPs, typically ∼10^4^ were expressed at a sufficient depth to be included in the analysis. Read-depth and alternate allele-fraction cutoffs were set at 30 and 0.05, respectively.

### Transgene Genotyping

The 5XFAD mouse is a model for familial Alzheimer’s disease that has overexpressing transgenes harboring mutations in *APP* (Amyloid Beta Precursor Protein) and *PSEN1* (Presenilin 1) that drive amyloid plaque formation.^6^ Transgene status, *i.e*. the presence of the 5XFAD transgenes, was established via two methods depending on the reference genome. If a sample was aligned to the mm39+5XFAD reference genome (n=164), read counts for 5XFAD-*APP* and 5XFAD-*PSEN1* were normalized to the library size, and the sample was determined to be transgenic if the normalized read counts were greater than 3×10^−6^ and 2×10^−6^, respectively. If a sample was aligned to the mm10 reference genome (n=215), unmapped reads were extracted with SAMtools and aligned to the 5XFAD sequence using BWA MEM.^13^ Read counts overlapping with 5XFAD coding regions were then normalized to the total library size and samples were called as transgenic if normalized read counts for 5XFAD-*APP* and 5XFAD-*PSEN1* were greater than 3×10^−5^ and 2×10^−5^, respectively. Transgene genotype validation was performed using DNA isolated from remaining tail tip tissue collected at time of harvest and processed according to the Jackson Laboratory Protocol 20498: QPCR Assay – Generic APP human genomic or cDNA, Version 5.0.

### Sex Matching

Sample sex was verified via the ratio of read-counts from *Xist*, a critical gene for the X-Inactivation process and the Y-chromosome gene *Ddx3y*. Samples with an *Xist*/*Ddx3y* ratio > 1 were designated as female, while samples with an *Xist*/*Ddx3y* ratio < 1 were designated as male. The presence of reads associated with *Ddx3y* typically occurs at low levels in female samples. This is likely due to the presence of cell-free RNA.

## Results

### Strain Matching

Of the 379 samples tested (294 B6xBXD and 85 CC), 332 (88%) returned the best matching score for the assigned strain (Table S1; mean score for assigned strain = 98% ± 3%) with the matching scores for all other strains ranging from 1-84% (mean = 51% ± 9%). The remaining 47 samples were comprised of 8 pairwise sample-swaps (16 mismatched samples), 1 false strain assignment, an offset in sample labels accounting for 15 mismatches, and one strip tube label swap (15 samples). In the tube swap, a 16^th^ sample was mismatched, but failed sequencing QC and was therefore not included in this analysis; additionally, independent of the tube swap, two samples among those swapped were also mislabeled. The distribution of matching scores by corrected sample strain can be found in Figure 1, with the correct strain in blue and all other strains in goldenrod.

**Figure 1.**
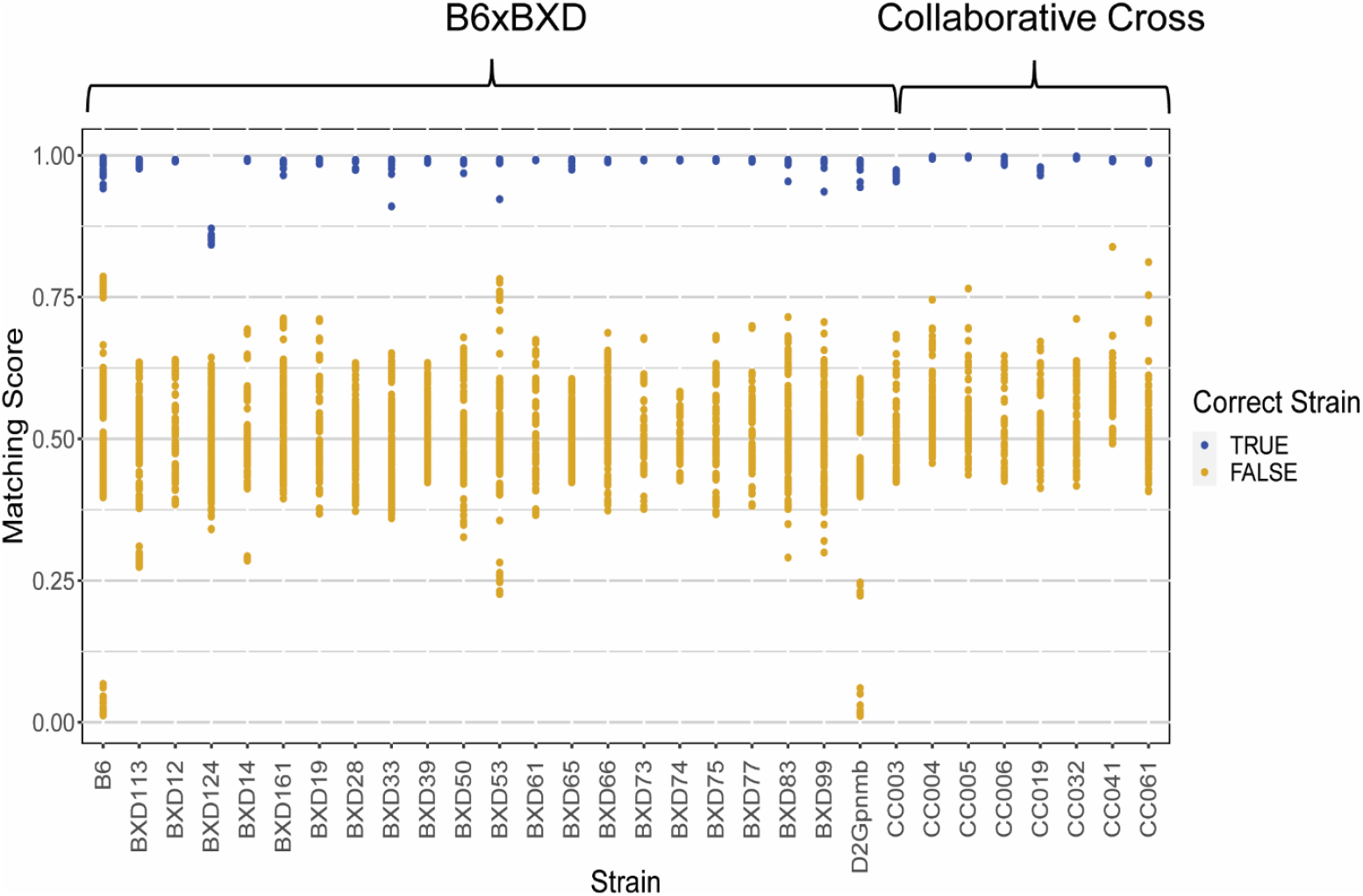
Matching scores by sample strain. Distributions of matching scores are plotted for each strain by correct sample strain (x-axis). The corresponding strain is shown in blue, while all other strains are shown in goldenrod.

**Figure 2.**
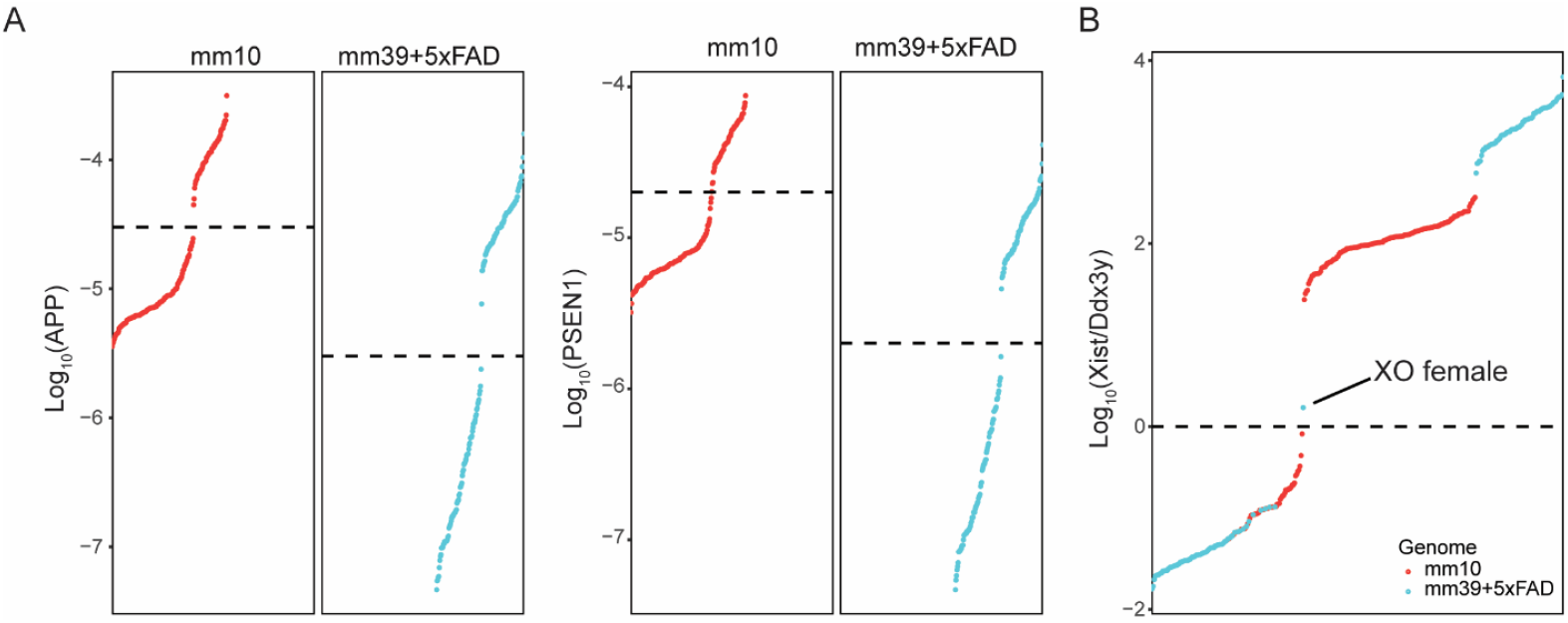
Base-10 log of reads associated with *APP* and *PSEN1* (a) and the *Xist*/*Ddx3y* ratio (b) ordered from lowest to highest values. Samples aligned to the mm10 genome are in red and samples aligned to the mm39+5XFAD genome are in blue. Dashed lines represent cutoffs used for non-transgenic vs. transgenic (a) and male vs. female (b), where non-transgenic and male samples land below the specified values.

The match rate among B6xBXD124 samples (mean = 85% ± 1%) was significantly lower than other strains (mean = 99% ± 1%; *t*-test *p*-value = 2.5e-21). All B6xBXD124 samples did, however, match BXD124 more closely than any other strain (Figure 1). This indicates that there may be significant disagreement between these genotypes and the genotypes of samples from which the NucSeq data was derived. We used the “varMatch” option in RNA Strain-Match to identify all SNPs that are mismatched to all samples in each B6xBXD strain for both the mm10- and mm39-aligned samples. Among the SNPs that were informative across B6xBXD124 samples, 11% were mismatched in mm10-aligned samples and 12% were mismatched in mm39-aligned samples. The average rate of consistently mismatched variants was 0.4% ± 0.2% for all other mm10-aligned strains and 0.4% ± 0.1% for all other mm39-aligned strains.

Notably, the lowest aligned match for B6 was D2, with average of 2.72% ± 1.52%, and vice versa it was 1.97% ± 1.28% for D2 matching with B6.

### Transgene Status

In looking at 5XFAD-*APP* and -*PSEN1* normalized read counts, we identified 27 samples that had mismatches between genomic and transcriptomic 5XFAD transgene status (transgenic/non-transgenic); 25 of these are attributed to sample mismatches identified by our strain analysis and were resolved upon correcting the sample-swaps and offset described above. Two additional samples were labelled as transgenic, but showed few reads associated with 5XFAD. These samples were taken from the HC and Ctx of the same mouse. qPCR-based genotype validation from genomic DNA extracted from tail tissue confirmed this sample as non-transgenic.

### Sex Matching

Using the *Xist*/*Ddx3y* ratio, 20 samples with mislabeled sex were identified; 19 of these were among the samples with mislabeled strains. One sample in the offset discussed above, however, was only identified by a mismatch in sex, *i.e*. the mismatch and the correctly identified sample shared both strain and transgene status. One additional sample (mouse ID: 1575) was identified as a female B6xD2Gpnmb mouse with a single X chromosome (XO). To provide further evidence that this was the case, we calculated the mean *D*-allele fractions on the X-chromosome and comparing to other B6xD2Gpnmb mice from the same study. In all cases, the *B*-allele is found on the maternal chromosome and the *D*-allele is found on the paternal chromosome; therefore, the *D*-allele should be largely absent from male and XO-female mice (non-zero *D*-allele fractions are expected in all samples due to cell-free RNA contamination as discussed above). The mean *D*-allele fraction on the X-chromosome for the suspected XO mouse was found to be 2.8e-3 ± 1e-2, which was comparable to those of the three male B6xD2Gpnmb mice from the same study (1.5e-3 ± 9e-3, 2.1e-3 ± 1e-2, and 2.0e-3 ± 1e-2) and much lower than that of the one XX-female B6xD2Gpnmb mouse from the same study (0.29 ± 0.2). Thus, this method can also be used to identify samples with spontaneous sex-chromosome aneuploidy.

## Discussion

For every sample that was tested using RNA Strain-Match, the sample’s correct strain was found to match its RNA data markedly better than all other tested strains. In the case of all transgene- and sex-mislabeled samples identified by RNA Strain-Match, we were able to identify the origin of the disparities and reassign the RNA data to its correct source. Had we not identified these mismatches, the corresponding studies would have been at risk of losing power in the best-case scenario, or presenting erroneous results in the worst-case scenario. This is particularly true in the case of Study 5 (Table 1), in which 41/159 (26%) samples were incorrectly assigned. Fortunately, in this case, no analyses had been conducted prior to performing these QC steps. For all other studies, analyses were redone with the corrected sample identification prior to publication.

This tool may be expanded to include other functionalities, such as alignment to protein data, and is part of our future directions.

## Summary

In our dataset of HC-, HT-, and Ctx-derived NucSeq samples from 294 B6xBXD and 85 CC mice, RNA Strain-Match successfully matched all samples to their appropriate strains. In combination with analyses of sample sex and transgene status, RNA Strain-Match identified a total of 50 mismatches. Of the mismatches identified, 40 were part of a single study that had a total of 159 samples. Had we not performed this QC step, >25% of the samples in that study would have been misrepresented, which would have dramatically impacted subsequent findings and interpretation of these data. Though we hope that this is an extreme example of sample misidentification, it effectively reenforces the importance of taking steps to verify that all samples are correctly labelled.

## Acknowledgments

We gratefully acknowledge the contribution of Drs. Bill Flynn, Eloise Courtois and the Single-Cell Biology Core Service at The Jackson Laboratory as well as the director of the computational sciences group at The Jackson Laboratory, Dr. Vivek Philip, for expert assistance with the work described in this publication. We are also thankful to funding from the National Institute of Aging: R01AG074012 (C.C.K.) and RF1AG059778 (C.C.K. and K.M.S.O).

## Figure and Table Legends

**Table S1**. Trial data used to test RNA Strain-Match. Strain, sex, and transgene matching data for each sample used in the study are included along with original and corrected metadata. Expected strain, transgene status, and sex are the values originally assigned to each sample prior to performing QC analysis. Expected strain score is the sample’s matching score associated with its expected strain. Strain, transgene status, and sex are the values assigned analytically using the metrics described in the methods section. Strain match, transgene status match, and sex match are Boolean values indicating if the associated feature matches its expected value.

## Notes

### Competing Interest Statement

The authors have declared no competing interest.

